## Supplementary figures and images for "The kinase Isr1 negatively regulates hexosamine biosynthesis in *S. cerevisiae*"

### S1 Fig

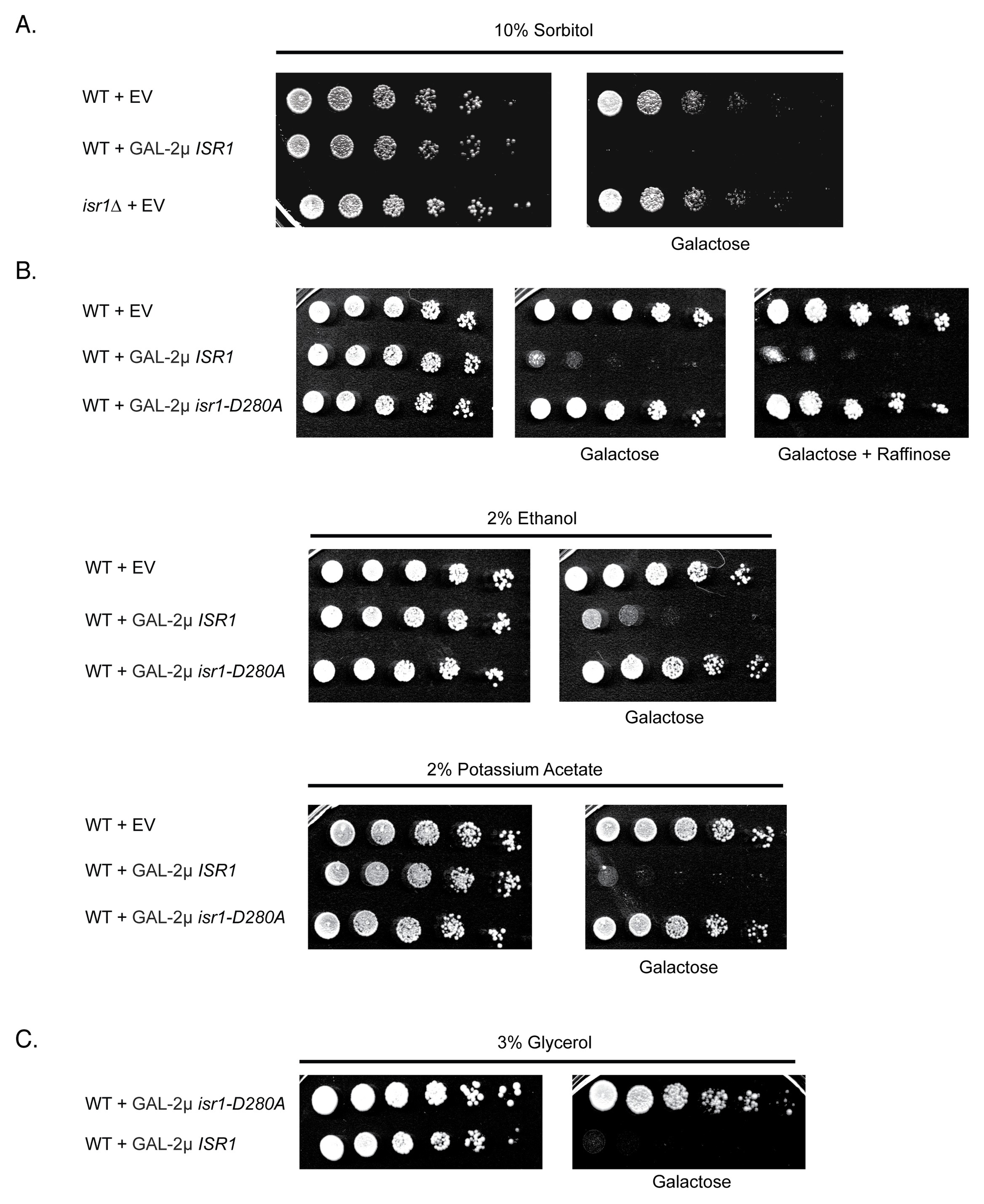

### S2 Fig

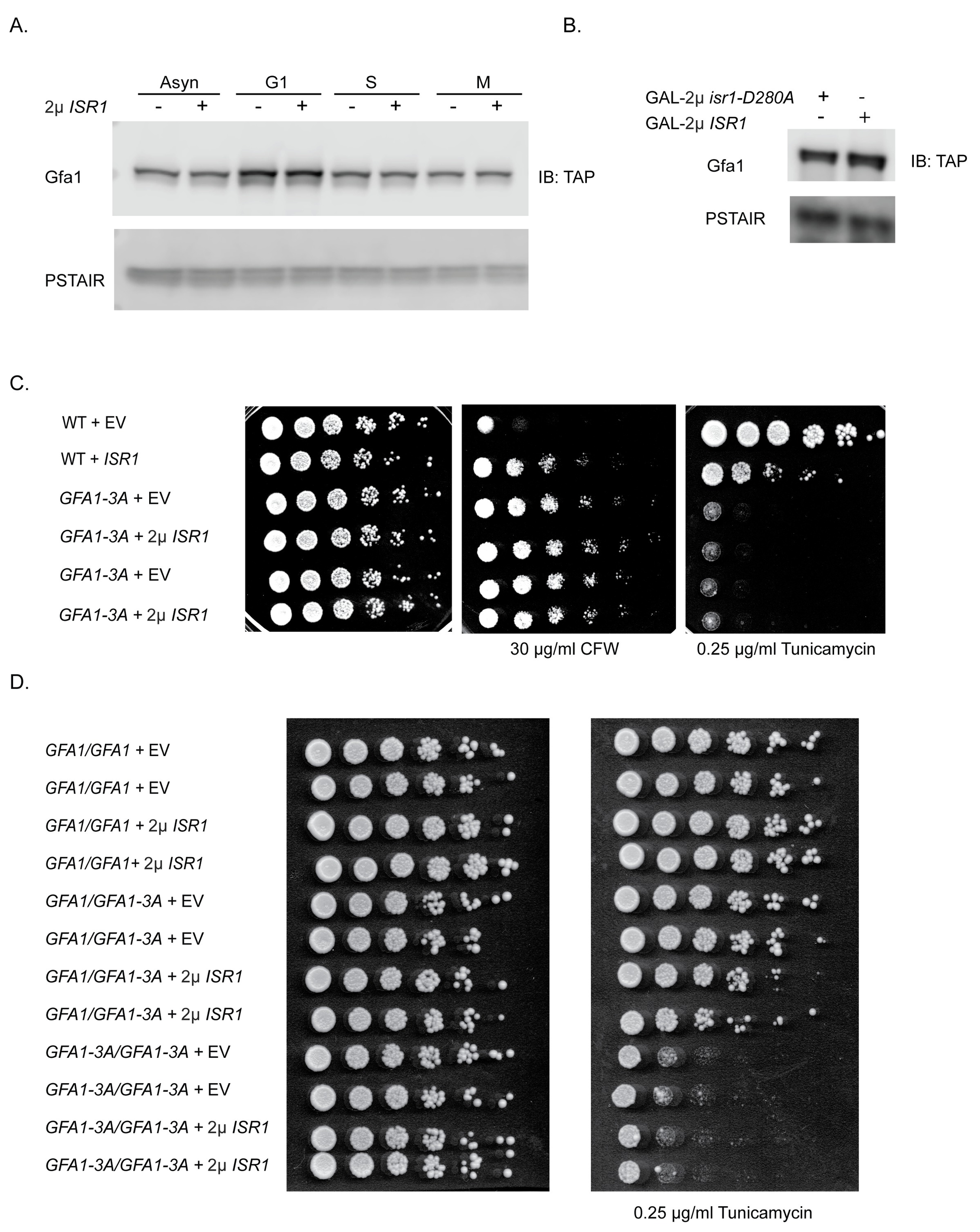

### S3 Fig

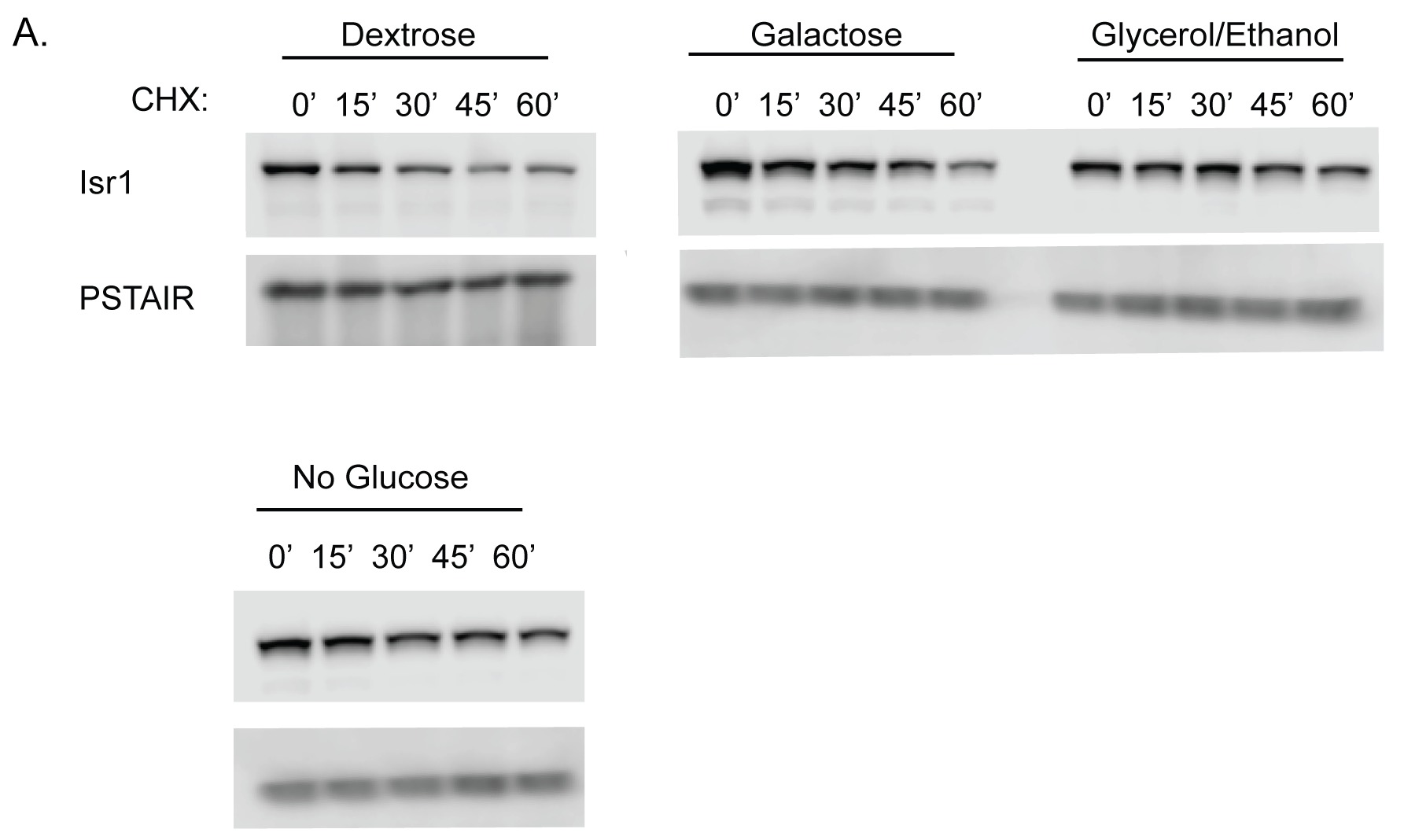

### S4 Fig

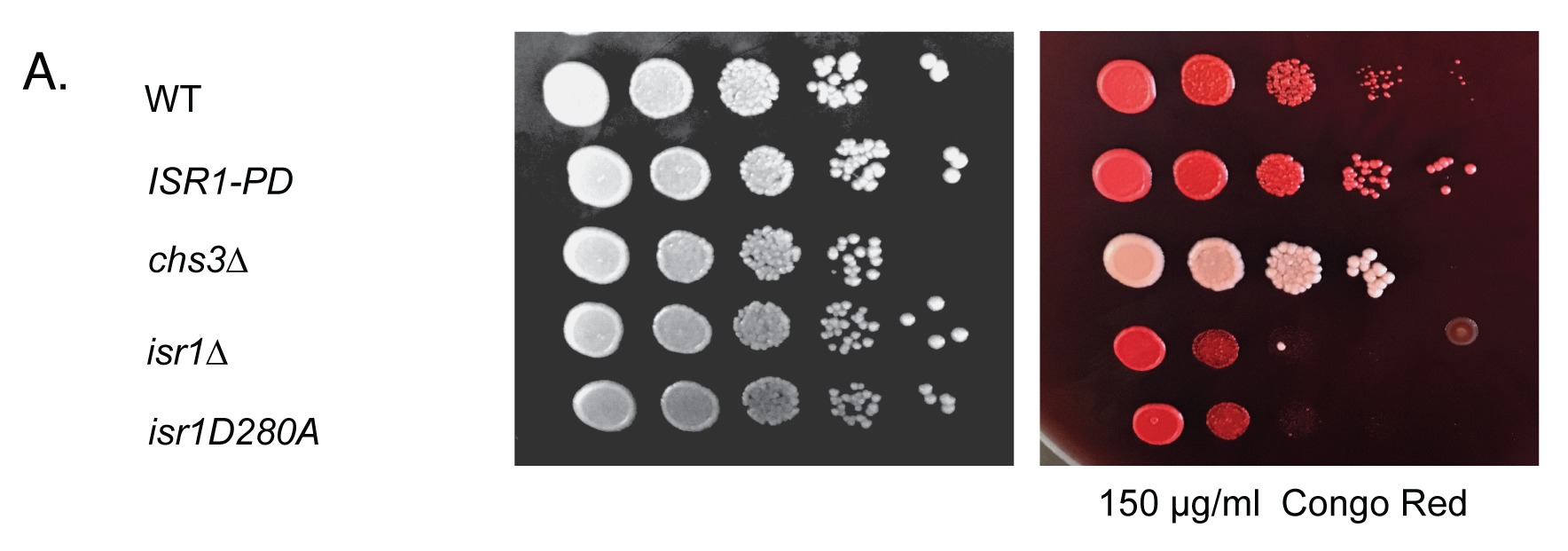

### S5 Fig

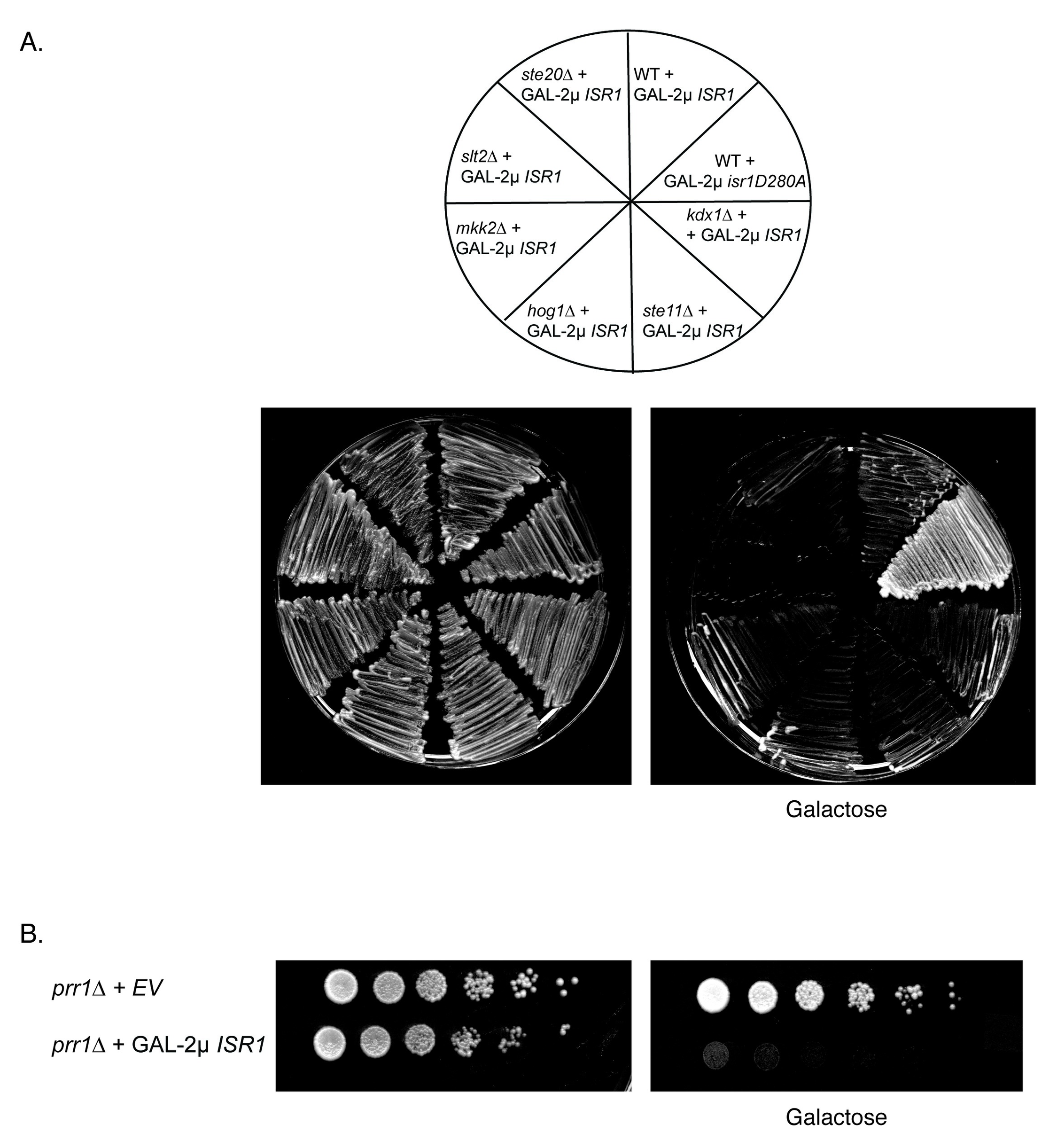
