## Supplementary material for "The kinase Isr1 negatively regulates hexosamine biosynthesis in *S. cerevisiae*": S6 Fig

**Fig S6.** ***ISR1-PD*** **sequence**

GTGATGTATGGGTCTATCTTATTGTTTATTTTGACTAGCATTGAATAAACAAAAAGC
GCTGCTAATAGATTCTTGCCTTTATATAACGGGGATCTAACCTCATCAACAGCAAAA
ATCGTCTTAAAACACATCTCAAAGACTAGTTCTCAAACGTCACGCTATGAACgctgCA
CCTCCTgccgCACCCGTCACCAGGGTTTCTGATGGTTCCTTTCCATCCATAAGTAAC
AATAGTAAGGGTTTTGCTTATCGCCAACCGCAAAAACATAAAAGTAACTTCGCATAT
TCACATCTGGTATCTCCTGTAGAGGAGCCGACAGCTAAATTCAGTGAGGCATTCCA
GACAGATTATTCTAGTAAGGCGCCCGTTGCTACCTCGGAGGCGCACCTAAAGAAC
GATTTAGACGTATTGTTCgCTgCCCCCCGGTTTTACgCTCCGGAGAATTTGGCTTTA
ATGTTCCGTCTTTCTAATACAGTTTCTTCCCTAGAATTTCTGGATGAGTTTTTGATGG
GCATATTACTTGCTCCAGAGATGGATTTTTTGTCAAATCCAAGTTATTCTCTTCCGT
CTAACAAATTAGTGGGACAGGGAAGTTATTCATATGTGTACCCTATATCATCAAGTG
CTTCATCAAGATGTAACAACGATTCAGGGGTTGTTTTAAAGTTTGCCAAATCACAGC
ATAAAAGCAAGGTGATTTTACAGGAAGCTTTGACGCTAGCATATCTCCAGTACATG
AGTCCTTCAAC
