## Supplementary material for "The kinase Isr1 negatively regulates hexosamine biosynthesis in *S. cerevisiae*": S2 Table

**Table S2. Strains used in this study**

| Strain Name | **Genotype** | **Fig** |
|---|---|---|
| BY4741 | *MATa his3Δ1 leu2Δ0 met15Δ0 ura3Δ0* | |
| EBA142 | *MATa his3Δ1 ura3Δ0 leu2Δ0 met15Δ0 isr1Δ::KanMX* | 2A |
| EBA211 | *MATa his3Δ1 ura3Δ0 leu2Δ0 met15Δ0 isr1Δ::HygMX* | 2A, 3C, 3D, 4B, 4C, 6C, 6D, S4 |
| EBA206 | *MATa his3Δ1 ura3Δ0 leu2Δ0 met15Δ0 Pkc1-3xFlag::KanMX* | 1C |
| EBA183 | *MATa his3Δ1 ura3Δ0 leu2Δ0 met15Δ0 ISR1-13xmyc::NAT* | 6C, 6D, S4 |
| EBA185 | *MATa his3Δ1 ura3Δ0 leu2Δ0 met15Δ0 ISR1-PD13xmyc::NAT* | 6C, 6D, 6E, S4A |
| EBA176 | *MATa his3Δ1 ura3Δ0 leu2Δ0 met15Δ0 isr1D280A::NAT* | 6C, 6D, S4 |
| EBA151 | *MATα his3Δ1 ura3Δ0 leu2Δ0 lys2Δ0 chs3Δ::KanMX* | 2B,6C, 6D, S4 |
| EBA160 | *MATa his3Δ1 ura3Δ0 leu2Δ0 met15Δ0 GFA1-TAP-His3MX* | S2 |
| EBA329 | *MATa his3Δ1 ura3Δ0 leu2Δ0 met15Δ0 GFA1-3A::HygMX* | 4D, 4E, S2 |
| EBA330 | *MATa his3Δ1 ura3Δ0 leu2Δ0 met15Δ0 GFA1-3A::HygMX* | 4D, 4E, S2 |
| EBA326 | *MATa/MATα his3Δ1/ his3Δ1 ura3Δ0/ ura3Δ0 leu2Δ0/ leu2Δ0 MET15/met15Δ0 LYS2/lys2Δ0 GFA1/GFA1-3A::HygMX* | 4F, S2 |
| EBA327 | *MATa/MATα his3Δ1/ his3Δ1 ura3Δ0/ ura3Δ0 leu2Δ0/ leu2Δ0 MET15/met15Δ0 LYS2/lys2Δ0 GFA1/GFA1-3A::HygMX* | 4F, S2 |
| EBA114 | *MATa/MATα his3Δ1/ his3Δ1 ura3Δ0/ ura3Δ0 leu2Δ0/ leu2Δ0 MET15/met15Δ0 LYS2/lys2Δ0* | 2E, 4F, 6E, S2 |
| EBA268 | *MATa/MATα his3Δ1/ his3Δ1 ura3Δ0/ ura3Δ0 leu2Δ0/ leu2Δ0 MET15/met15Δ0 LYS2/lys2Δ0 ISR1/ISR1-PD::NAT* | 6E |
| EBA273 | *MATa/MATα his3Δ1/ his3Δ1 ura3Δ0/ ura3Δ0 leu2Δ0/ leu2Δ0 MET15/met15Δ0 LYS2/lys2Δ0 GFA1/gfa1Δ::KanMX* | 2E, 6E, S2 |
| EBA269 | *MATa/MATα his3Δ1/ his3Δ1 ura3Δ0/ ura3Δ0 leu2Δ0/ leu2Δ0 MET15/met15Δ0 LYS2/lys2Δ0 GFA1/gfa1Δ::KanMX ISR1/ISR1-PD::NAT* | 6E |

| | | |
|---|---|---|
| EBA368 | *MATa/MATα his3Δ1/ his3Δ1 ura3Δ0/ ura3Δ0 leu2Δ0/ leu2Δ0 MET15/met15Δ0 LYS2/lys2∆0 GFA1-3A/GFA1-3A::HygMX* | S2 |
| EBA369 | *MATa/MATα his3Δ1/ his3Δ1 ura3Δ0/ ura3Δ0 leu2Δ0/ leu2Δ0 MET15/met15Δ0 LYS2/lys2∆0 GFA1-3A/GFA1-3A::HygMX* | S2 |
| EBA315 | *MATa/MATα his3Δ1/ his3Δ1 ura3Δ0/ ura3Δ0 leu2Δ0/ leu2Δ0 MET15/met15Δ0 LYS2/lys2∆0 GNA1/gna1∆::KANMX ISR1/ISR1-PD::NAT* | 6E |
| EBA316 | *MATa/MATα his3Δ1/ his3Δ1 ura3Δ0/ ura3Δ0 leu2Δ0/ leu2Δ0 MET15/met15Δ0 LYS2/lys2∆0 GNA1/gna1∆::∆::KANMX* | 2E, 6E |
| EBA302 | *MATa/MATα his3Δ1/ his3Δ1 ura3Δ0/ ura3Δ0 leu2Δ0/ leu2Δ0 MET15/met15Δ0 LYS2/lys2∆0 GLN1/gln1∆::KANMX ISR1/ISR1-PD::NAT* | 6E |
| EBA319 | *MATa/MATα his3Δ1/ his3Δ1 ura3Δ0/ ura3Δ0 leu2Δ0/ leu2Δ0 MET15/met15Δ0 LYS2/lys2∆0 PCM1/pcm1∆::KANMX* | 2E, 6E |
| EBA328 | *MATa/MATα his3Δ1/ his3Δ1 ura3Δ0/ ura3Δ0 leu2Δ0/ leu2Δ0 MET15/met15Δ0 LYS2/lys2∆0 PCM1/pcm1∆::KANMX ISR1/ISR1-PD::NAT* | 6E |
| EBA304 | *MATa/MATα his3Δ1/ his3Δ1 ura3Δ0/ ura3Δ0 leu2Δ0/ leu2Δ0 MET15/met15Δ0 LYS2/lys2∆0 QRi1/qri1∆::KANMX* | 2E, 6E |
| EBA303 | *MATa/MATα his3Δ1/ his3Δ1 ura3Δ0/ ura3Δ0 leu2Δ0/ leu2Δ0 MET15/met15Δ0 LYS2/lys2∆0 QRi1/qri1∆∆::KANMX ISR1/ISR1-PD::NAT* | 6E |
| EBA135 | *MATa his3Δ1 ura3Δ0 leu2Δ0 met15Δ0 Isr1-13xmyc::URA3* | 5A, 5B, 5C, 5D, 5E, 6B, S3 |
| EBA153 | MATa his3Δ1 ura3Δ0 leu2Δ0 met15Δ0 pho85∆::KANMX Isr1-13xmyc::URA3 | 5C, 6B |
| EBA332 | MATa his3Δ1 ura3Δ0 leu2Δ0 met15Δ0 cdc4-1::HYGMX Isr1-13xmyc::URA3 | 5B |
| EBA158 | MATα his3Δ1 ura3Δ0 leu2Δ0 lys2∆0 cdc53-1 Isr1-13xmyc::URA3 | 5B |
| EBA331 | MATa his3Δ1 ura3Δ0 leu2Δ0 met15Δ0 pcl1∆::KANMX Isr1-13xmyc::URA3 | 5C |
| EBA174 | MATa his3Δ1 ura3Δ0 leu2Δ0 met15Δ0 3xHA-Isr1∆93::NATMX | 6D |

| | |
|---|---|
| knockout collection | MATa his3Δ1 ura3Δ0 leu2Δ0 met15Δ0 YFGΔ::KANMX |
