## Supplementary material for "The kinase Isr1 negatively regulates hexosamine biosynthesis in *S. cerevisiae*": S3 Table

**Table S3: Plasmids used in this study**

| Plasmid Name | Description | Fig/Use |
|---|---|---|
| EBP294 | Prs426 NAT (EV) | 1A, 2B, 3B, 3C, 3D, 4D, 4E, S1, S2, S5 |
| EBP187 | 2µ *ISR1* | 1B, 2B, 3C, 4E, S2 |
| EBP290 | 2µ *ISR1 isr1-D280A* | 2B, 3C |
| EBP211 | Gal- 2µ *ISR1* | 1, 2C, 2E, 3B, 3D, 4A, 4B, 4C, 4D,4F, S1, S2, S5 |
| EBP210 | Gal- 2µ *isr1-D280A* | 1A, 2C, 3B, 3D, 4A, S1, S2 |
| EBP215 | prs426NAT + *GAL1-ISR13xFlag*, *GFA1pr-GFA1* | 4A |
| EBP216 | pr*s426NAT* + *GAL1-isr1D280A-3xFlag, GFA1pr-GFA1* | 4A |
| EBP211 | prs426NAT + *GAL1-ISR1-3xFlag*, *QRI1pr-QRI1* | 4A |
| EBP212 | prs426NAT + *GAL1-isr1D280A-3xFlag*, *QRI1pr-QRI1* | 4A |
| EBP168 | prs402NAT + *ISR1pr-ISR1-13xMyc* | Construction of *ISR1* integration (wildtype) |
| EBP181 | prs402NAT + *ISR1pr-ISR1PD-13xMyc* | Construction of *ISR1-PD* integration |
| EBP172 | prs402NAT + *ISR1pr-isr1D280A* | Construction of *isr1-D280A* integration |

| | | |
|---|---|---|
| EBP220 | p3xFlagHYGMX + *GFA1* | Construction of *GFA1* integration (WT) |
| EBP223 | p3xFlagHYGMX + *GFA1-S332AT334AS336A* | Construction of *GFA1-3A* integration |
| MS197 | Prs306 + *ISR1-13xMyc* | Construction of *ISR1-13xmyc* tagged strains |
| PYMN-20 | PYMN-20 (66) | Construction of *HA-ISR1Δ93* |
